# Widespread polycistronic transcripts in mushroom-forming fungi revealed by single-molecule long-read mRNA sequencing

**DOI:** 10.1101/012542

**Authors:** Sean P. Gordon, Elizabeth Tseng, Asaf Salamov, Jiwei Zhang, Xiandong Meng, Zhiying Zhao, Dongwan Kang, Jason Underwood, Igor V. Grigoriev, Melania Figueroa, Jonathan S. Schilling, Feng Chen, Zhong Wang

## Abstract

Genes in prokaryotic genomes are often arranged into clusters and co-transcribed into polycistronic RNAs. Isolated examples of polycistronic RNAs were also reported in some eukaryotes but their presence was generally considered rare. Here we developed a long-read sequencing strategy to identify polycistronic transcripts in several mushroom forming fungal species including *Plicaturopsis crispa, Phanerochaete chrysosporium, Trametes versicolor* and Gloeophyllum trabeum^1^. We found genome-wide prevalence of polycistronic transcription in these Agaricomycetes, and it involves up to 8% of the transcribed genes. Unlike polycistronic mRNAs in prokaryotes, these co-transcribed genes are also independently transcribed, and upstream transcription may interfere downstream transcription. Further comparative genomic analysis indicates that polycistronic transcription is likely a feature unique to these fungi. In addition, we also systematically demonstrated that short-read assembly is insufficient for mRNA isoform discovery, especially for isoform-rich loci. In summary, our study revealed, for the first time, the genome prevalence of polycistronic transcription in a subset of fungi. Futhermore, our long-read sequencing approach combined with bioinformatics pipeline is a generic powerful tool for precise characterization of complex transcriptomes.

## Significance Statement

Our long-read sequencing led to an unexpected discovery of the genome-wide presence of polycistronic transcripts in mushroom forming fungi. In contrast to the previous belief that polycistronic transcription is largely restricted to prokaryotes, we demonstrated it is also a conserved feature unique to mushroom forming fungi, with a potential role in regulating gene expression. This study, for the first time, suggests genome-wide polycistronic transcription may not be a unique feature to prokaryotes.

## Introduction

Advances in sequencing technologies have led to the discovery of an enormous variety of RNA species within cells, including both coding and non-coding RNAs^2,3^, splicing isoforms^4^, alternatively polyadenylated isoforms^5,6^, and gene-fusion transcripts^7-9^. High-throughput short-read sequencing of transcriptomes (RNA-seq) has enabled a precise quantification of gene expression levels and the identification of new exons and splice junctions^10,11^. However, as short-reads are much shorter than the length of most transcripts, assembly of these short-reads is necessary to infer the full cornucopia of transcript diversity^12^. For organisms that lack a reference genome, *de novo* transcriptome assembly from short-reads is often the only available choice. However, transcript assembly has many informatics challenges as it involves piecing together large volumes of short-reads to reconstruct individual transcript isoforms^12^. The largest challenges of short-read assembly include resolving hundreds of distinct isoforms derived from the same loci, and overlapping transcripts on the same strand for transcripts that span different loci^9, 13, 14^. Reduced sensitivity of short-read assembly to identify multiple isoforms from the same locus and long multi-locus transcripts clouds our ability to accurately define transcriptional units.

With a mean read length of ∼7 kb, the Pacific Biosciences (PacBio) single-molecule sequencing platform provides a direct and unbiased observation of full-length transcripts and their diversity. The throughput of the technology has dramatically increased, making genome-wide transcriptome studies possible for eukaryotes^15-17^. To overcome the low single-pass sequencing accuracy of the platform, recent studies either used circular consensus (CCS) reads^16^ or 2^nd^ generation short-reads to correct errors in PacBio long-reads^17, 18^. The CCS correction strategy excludes long transcripts (>3kb) and thus has limited ability to analyze long RNAs, while the short-read correction strategy requires additional sequencing efforts and the short-read sequencing may have biased coverage over transcripts with extreme GC-content. Thus, additional approaches are needed to fully utilize PacBio long-reads for comprehensive transcriptomics studies.

In this study we developed a transcriptome sequencing and analysis strategy called ToFU (<u>T</u>ranscript is<u>O</u>forms: <u>F</u>ull-length and <u>U</u>nassembled) that requires only PacBio reads for generating a *de novo* transcriptome, eliminating the need for short-read assembly or reference genomes. We chose to test ToFU on four wood-degrading basidiomycete fungal transcriptomes, *Plicaturopsis crispa, Phanerochaete chrysosporium, Trametes versicolor* and *Gloeophyllum trabeum* as these fungi possess genomic characteristics that are ideal to examine the effectiveness of our approach. First, these basidiomycete fungi have genes with higher intron numbers and more prevalent alternative splicing than ascomycetes and thus rich RNA isoform diversity^19^. Second, despite exhibiting complex alternative splicing, intron-rich basidiomycetes have smaller numbers of expressed loci than many higher eukaryotes, which makes them an ideal candidate for testing ToFU. Finally, the biochemical and physiological adaptations of these fungi to decompose wood represent a mechanism with great biotechnological potential in engineering plant biomass deconstruction and advancing synthetic biology. Our knowledge related to RNA transcript isoform diversity in intron-rich fungi is limited as they are under-represented in transcriptome studies, and little is known about isoform diversity of mRNAs encoding the enzymes that govern wood-degrading processes. With these in mind, we first deeply sequenced the transcriptome of the white-rot basidiomycete *P. crispa* with both short- and long-read technology to benchmark our approach. Subsequently, we generated additional long-read transcript sequences for three additional species (*P. chrysosporium, T. versicolor* and *G. trabeum*) representing different orders within the Basidiomycota and showed the existence of widespread long polycistronic mRNAs in mushroom-forming fungi.

## Results

### A single-molecule, long-read strategy to identify full-length isoforms

The goal of ToFU was to bypass complicated experimental and informatic procedures of short-read assembly and instead leverage the longest reads from the PacBio platform to yield high-confidence transcript isoforms independent of a reference genome and therefore making the approach applicable to any organism (Figure 1 and Methods). To increase the representation of different mRNA populations in *P. crispa*, multiple cDNA libraries, including size selected (1-2 kb, 2-3 kb, and 3-6 kb) and non-size selected libraries, were generated and sequenced for each of two growth conditions. After sequencing, we identified putative full-length cDNA reads from 5 million raw reads by the presence of both 5’ cDNA primers and polyA signals preceding the 3’ primers, yielding 2.1 million full-length sequences. Reads derived from the same isoforms were then clustered to generate initial consensus sequences, and further polished with the aid of non-full-length reads to generate 176,903 high-quality consensus sequences. After merging redundant sequences we obtained 22,956 distinct isoforms representing 9,073 transcribed loci (Supplemental Table 1). In the following sections, we denote this final set of isoforms as the ToFU transcript set. For performance comparison and validation purposes, we also generated 300 million paired-end 100bp short-reads on the Illumina HiSeq platform from the same RNA samples.

**Figure 1.**
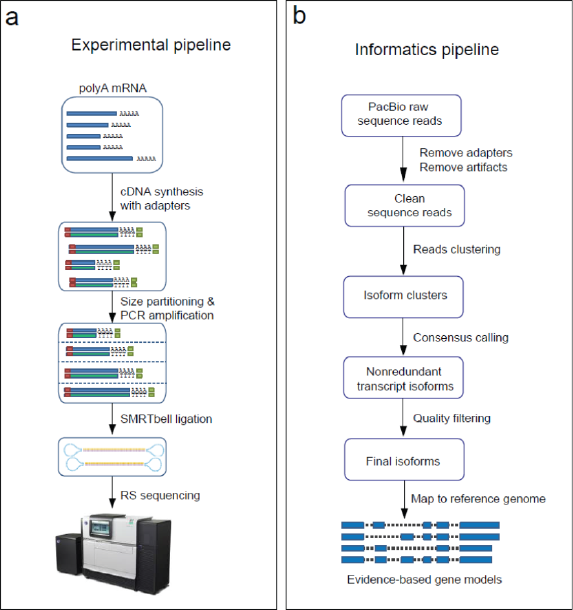
An overview of the experimental (a) and informatics (b) components in the ToFU pipeline to generate transcript isoforms.

### ToFU transcripts are long and accurate

The ToFU transcripts (Fig 2a) have an average length of 1,657 nt, with the longest being 5,589 nt. The length of the final transcripts closely follows the distribution of the input full-length reads (*Input FL Reads* in Fig 2a) since no assembly is involved. ToFU transcripts include a large number of isoforms greater than 3 kb that are not accessible by simply using CCS reads (*HQ CCS Reads*, Fig 2a)^16^.

**Figure 2.**
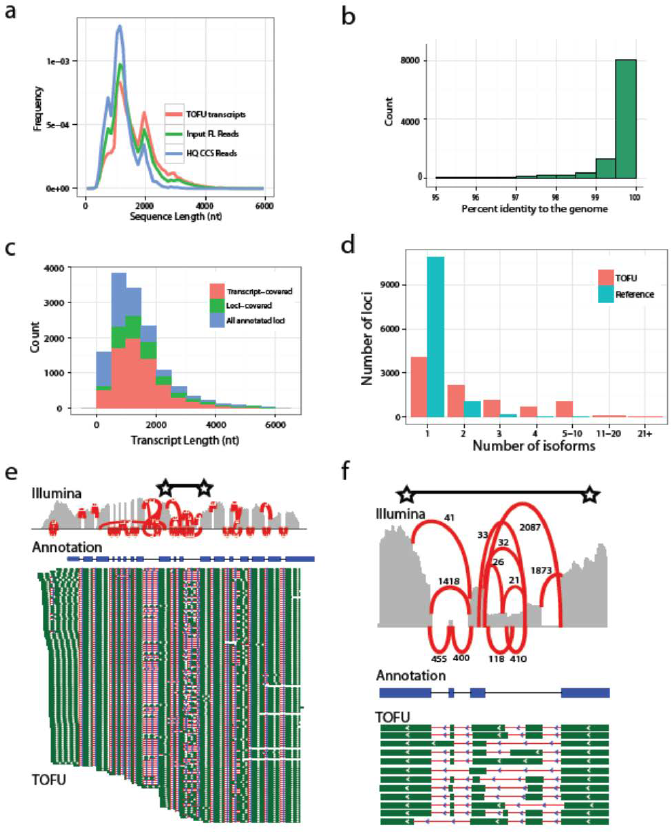
Long, high-quality, consensus sequences accurately benchmark transcript diversity. **a**, Length distributions of full-length (FL) input reads, high-quality CCS reads, and ToFU transcript sequences. **b**, Histogram of percent nucleotide identity of ToFU transcript sequences aligned to the reference genome. **c**, Accumulative histogram of number of reference annotations that have a ToFU transcript that completely covers each annotated junction (transcript-covered) or only partially covers the loci (loci-covered). **d**, Distribution of distinct isoforms per loci for the reference annotation and ToFU transcript set. **e**. Illumina short-read coverage (grey) and junction support (red) aligned along the reference annotated transcript (blue) for a glycosyl hydrolase gene with 120 distinct PacBio isoforms aligned below (splice junctions are shown in red and exon sequences are shown in green). **f**, An enlarged view of the region between two starts in **2e**.

Despite the ∼15% error rate in the input reads^18^, our analyses indicate that ToFU transcripts are highly accurate. Although our pipeline did not require a sequenced genome, we used the annotated draft genome sequence of *P. crispa* from JGI MycoCosm portal^20^ (http://jgi.doe.gov/Plicaturopsis) to independently estimate transcript accuracy. When aligned to the genome sequence using GMAP^21^ and allowed alignment gaps, 99.79% (37,930,451/38,011,774) of the bases are concordant with the reference base (Fig. 2b). The estimated errors for substitution, insertion, and deletion are 0.06%, 0.04% and 0.12% respectively. These percentages are likely over-estimated since they do not account for errors in draft reference genome, polymorphisms, or post-transcriptional RNA-editing. In addition, based on existing reference-based gene annotations^20^ the ToFU transcripts fully span most of the genes with detected expression (Figure 2c) and on average they have longer untranslated regions (UTRs) (Supplementary Fig. 1).

### ToFU reveals extensive alternative splicing and alternative poly-adenylation

Fungal species were previously thought to have much lower rates of alternative splicing than plants and animals. Recent estimates based on EST and RNA-Seq data suggest that on average approximately 7.3% of genes in non-Saccharomycotina fungi undergo AS, with *Cryptococcus neoformans* being an extreme case with up to 20% of genes involved in AS^19^. By contrast, 42% of genes in Arabidopsis and 95% in humans are alternatively spliced^22, 23^. Among 9,073 transcribed loci in *P. cripsa*, 56% (5,038 / 9,073) have two or more and 32% (2,908) have three or more distinct isoforms that derived from either alternative splicing, alternative poly-adenylation, or alternative transcription start sites (Fig. 2d). In total, 25.2% of all transcribed loci are alternatively spliced and 28.7% loci have alternative poly-adenylation sites. This estimation of splicing rate is likely underestimated, as rare isoforms may skip detection, and we only sampled two conditions. These findings suggest that basidiomycete fungi may have a much higher transcriptional diversity than previously reported.

Wood-decaying fungi produce a wide range of enzymes to break down plant cell walls including a large and diverse family of glycosyl hydrolases (GHs). Despite their importance, little is known about GH transcript isoform diversity at individual genes, which may affect the efficiency at which these enzymes are made both in their native host and bioengineered systems. Interestingly, among 151 loci that have 10 or more isoforms, 8 are associated with GH activity. One of these GH loci produces 120 distinct isoforms, with additional support from short-read validation of individual splice junctions (Fig. 2e,f).

### A quality evaluation of short-read assemblers

Extensive alternative splicing in the *P. crispa* transcriptome makes it a good candidate to assess the quality of algorithms for transcriptome reconstruction from short reads. In order to quantify the ability of existing short-read transcript reconstruction methods to capture isoform level resolution we used the ToFU transcript set as a reference. We selected three assemblers to represent both genome-based (Cufflinks^24^) and *de novo* (Rnnotator^25^ and Oases^26^) reconstruction strategies. All assemblies were generated from the above 300 million 100-bp paired end short-read dataset. The performance of each assembler was evaluated by its ability to recover ToFU transcripts (sensitivity) and the number of predictions validated by ToFU (specificity) (Figure 3). For a fair comparison, we only considered loci that were detected by both short-reads and ToFU transcripts, and we evaluated the reconstructed transcripts only based on their exon structures (splicing junctions).

**Figure 3.**
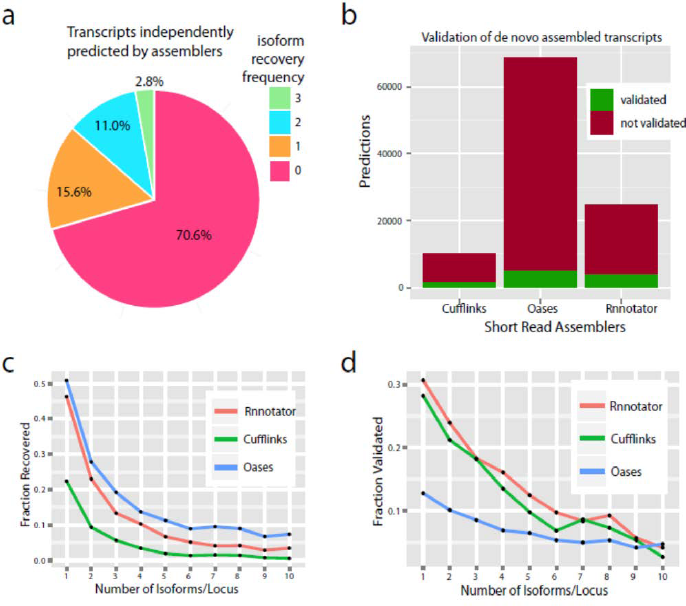
Evaluating short-read transcript reconstruction against ToFU transcripts. **a**, Percentage of ToFU transcripts recovered by three different short-read assembly methods. The isoform frequency shows whether a ToFU transcript is recovered by exactly 0, 1, 2, or all 3 of the assemblers. **b**, Number of assembled transcripts validated by ToFU transcripts. A transcript is validated as an exact match of a ToFU transcript if it shares exactly the same number of exons and donor-acceptor sites. **c**, Fraction of ToFU transcripts recovered (sensitivity) by each short-read assembler as a function of isoform complexity. **d**, Fraction of assembled transcripts validated (specificity) by ToFU as a function of isoform complexity. Isoform complexity is determined by the number of ToFU isoforms at each locus.

Overall, a single assembler was only able to reconstruct a small percentage of 22,956 ToFU isoforms, and only 2.8% of isoforms by all three methods (Fig. 3a). 70% of ToFU transcripts were not fully reconstructed by any of the three assemblers. Among the three short-read assemblers, Oases had the largest number of transcripts and the highest prediction sensitivity, but it also had the most predictions not validated by ToFU and thus the least specificity. Cufflinks seemed to be the most conservative assembler, predicting only a small number of transcripts compared with the other two. Rnnotator showed a balance between sensitivity and specificity, with only one-third as many transcripts predicted as Oases with similar sensitivity (Supplemental Tables 2, 3 and Fig 3b). Importantly, both the sensitivity (Fig 3c) and specificity (Fig 3d) of all the above assemblers dropped sharply as isoform complexity increased.

The above analyses highlight the limitations of current state-of-the-art short-read assembly methods for isoform discovery, and suggest that long-read RNA sequencing is essential for accurate isoform resolution, especially for genes with many isoforms.

### Long-read sequencing reveals widespread polycistronic mRNAs in *P. crispa*

Detailed analysis of the opening reading frames (ORFs) of the *P. crispa* ToFU transcript set revealed 508 genomic locations that produce readthrough mRNAs composed of two to four ORFs from either the same or opposite strands (an example is shown in Figure 4a). Three-hundred and fourteen (61.8%) of these readthrough transcripts containing two or more annotated reference genes were characterized further. They collectively involve 717 of the 9,073 transcribed loci (7.9%). Unlike the small regulatory upstream ORFs found in yeast and many other organisms^10, 27^, the average size of the upstream ORFs is comparable to the downstream ones (256 vs 277 amino acids) with an average inter-ORF distance of 364 nt.

**Figure 4.**
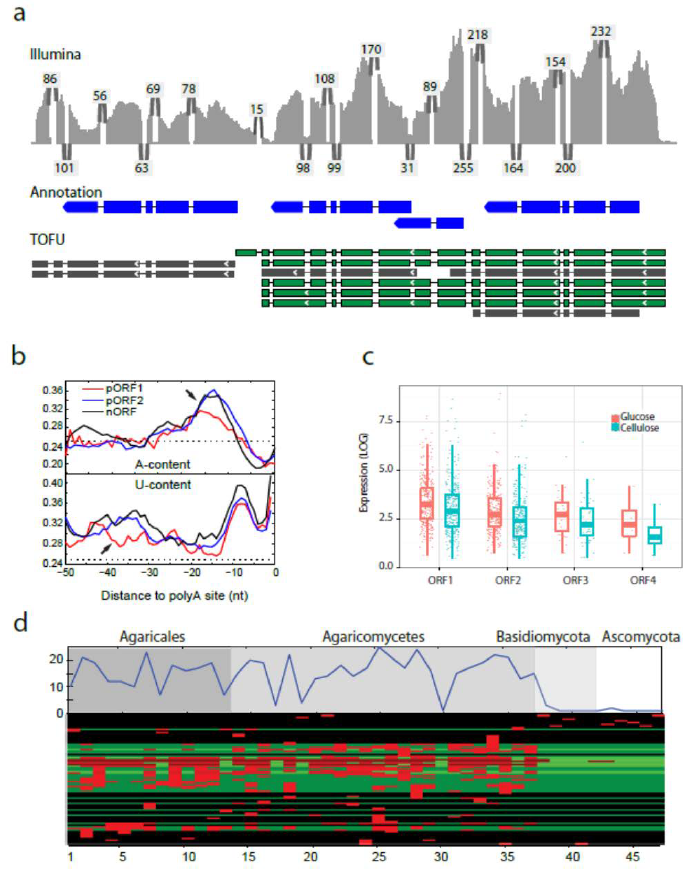
The genome-wide presence of polycistronic mRNAs. **a**, Short-reads (Illumina) aligned to a cluster of tandem reference genes (Annotation, 3 tandem genes on the first row). The numbers of supporting short-reads for each junction are indicated. Polycistronic transcripts (TOFU) are shown in green and non-polycistronic transcripts in gray. **b**, A comparison of transcription termination signals. The sequence composition profiles (upper panel for A-content and lower panel for U-content) before the polyadenylation sites for different classes of ORFs. pORF1 is the upstream ORF and pORF2 is downstream ORF, while nORF stands for non-polycistronic mRNAs. The y-axis are the frequencies of a specific nucleotide averaged for 200 randomly sampled polycistronic mRNA or non-polycistronic controls, dotted lines are the expected frequencies (0.25) if all four bases are equally likely. Arrows denote NUE (upper panel) and FUE (lower panel), respectively. For this figure, only polycistronic transcripts with exactly two ORFs are plotted. Genome-wide analysis base composition of termination signals for all transcribed loci is shown in Supplementary Fig. 2. **c**, The independent expression levels of ORFs within polycistronic RNAs. ORF numbers indicate their order in the transcript (5’- to 3’). **d**, Polycistronic transcripts are likely a unique feature to Agaricomycetes. The top plot shows the total number of adjacent ORF pairs within polycistronic transcripts from *P. crispa* that have conserved gene configuration in related species. The numbers on x-axis are species with increasing evolutionary distance. The bottom heatmap shows the conservation for each individual pair of ORFs. Red indicates the presence of a homologous gene pair in the species.

In the majority of cases (73%), ORFs within a readthrough transcript are in the same forward orientation; thus these transcripts are polycistronic mRNAs. Multiple stop codons are present in all reading frames between most of these ORFs, excluding the possibility that the transcripts are large single ORFs that are misannotated. Polycistronic transcripts are a common feature of the prokaryotes, but are relatively rare in eukaryotes except for transpliced transcripts in protists and nematodes^28^. To our knowledge this is the first report of polycistronic transcription in higher fungi. To rule out the possibility that these transcripts are experimental or informatics artifacts, we carried out independent validation experiments by RT-PCR followed by additional sequencing of amplicons (Methods and Supplemental Table 4). In support of the high fidelity of long-read sequencing strategy, 8 out of 10 randomly selected polycistronic transcripts were successfully validated, while the remaining 2 were inconclusive due to technical PCR problems.

In humans, *Arabidopsis thaliana* and the filamentous ascomycete *Aspergillus oryzae*, signals for transcription termination include an A-rich near upstream element (NUE), and a U-rich far upstream element (FUE)^5, 29, 30^. The polycistronic transcripts could result from transcriptional readthrough due to weak termination signals^5^. To address this possibility, we compared the sequence composition before the poly-adenylation sites of the upstream and the downstream ORFs. First, we did a genome-wide analysis of sequences surrounding poly-adenylation sites to confirm the presence of NUE and FUE elements in the basidiomycete, *P. crispa* (Supplementary Fig. 2). We then compared the termination signals of ORF1 in polycistronic transcripts (Figure 4b, pORF1) against its downstream ORF (Figure 4b, pORF2) and non-polycistronic transcripts (Figure 4b, nORF). Consistent with the weak transcription termination hypothesis, ORF1 is lacking both the U-rich FUE and A-rich NUE.

### Genes within polycistronic transcripts are also independently transcribed

Unlike in prokaryotes where polycistronic genes are transcribed into a single transcript without independent transcription, the polycistronic genes in *P. crispa* are also independently transcribed (Figure 4a). However, expression of downstream genes is consistently lower than their upstream counterparts within the same polycistronic transcript and this trend was consistent in independent experimental conditions (Figure 4c). Thus genes associated with readthrough transcripts frequently formed 2 to 4 successive tiers of decreasing gene expression. Polycistronic readthrough RNAs associated with this biased expression are different than previously identified regulatory RNAs such as lncRNAs^2^ that fall largely outside genic regions or transcripts that are found in antisense orientation relative to genes due to convergent readthrough transcription^31, 32^. This raises the possibility that the expression of downstream genes is repressed by the upstream readthrough transcription either through transcriptional interference (TI) or nucleosome positioning in which intergenic transcription alters the organization of nucleosomes at promoters thus influencing their activity^33, 34^.

### Polycistronic RNAs are likely a feature unique to Agaricomycetes

To investigate the evolutionary origin of these genome-wide polycistronic transcripts from *P. crispa*, we used the pairs of adjacent ORFs within these transcripts as queries to search 47 sequenced fungal genomes (Supplemental Table 5) for conserved gene configurations. These fungal species include 13 species from the same subclass as P. crispa (Agaricomycetidae), 24 from the same class (Agaricomycotina), and 33 from the same phylum (Basidiomycota). In addition, there are 4 species from the phylum Ascomycota. Since there are no available long read sequences from these species, we reasoned that conserved gene configuration would be indicative of possible readthrough transcription in other species. These conservation analyses indicate that only a subset of the gene pairs have conserved configuration in multiple species (Figure 4d), and this conservation declines sharply outside of the Agaricomycete class. This suggests that the gene pairs producing polycistronic transcripts in P. crispa may also produce polycistronic transcripts in other Agaricomycetes. This also implies that either polycistronic transcription is unique to Agaricomycetes or that more distally related species produce polycistronic transcripts from different pairs of adjacent genes.

To validate the notion that other Agaricomycetes produce polycistronic transcripts, we generated additional PacBio transcriptome data from three species from different orders than *P. crispa*, but within Agaricomycetes class. These fungi represent three independent orders (Polyporales, Gloeophyllales, Amylocorticiales), and include two additional white rot fungi *Phanerochaete chrysosporium*, and *Trametes versicolor*, as well as one brown rot fungus *Gloeophyllum trabeum* (Methods). Even without deep sequencing, we identified at least a hundred putative polycistronic readthrough transcripts from each fungus (Table 1). Among the *P. crispa* polycistronic gene pairs with homologous gene configurations in these species, PacBio long reads confirmed polycistronic transcripts associated with three gene pairs in *Trametes versicolor*, four gene pairs in *Phanerochaete chrysosporium* and one gene pair in *Gloeophyllum trabeum* (19, 19 and 21 total conserved gene pairs per species, respectively, Supplemental Table S5). To provide support for the absence of polycistronic transcription in non-Agaricomycetes we analyzed a deep long-read transcriptome data set for the ascomycete *Neurospora crassa*^35^ (*N. crassa*). This analysis did not identify any high confidence polycistronic transcripts. These results are consistent with the above hypothesis that genome-wide polycistronic transcription is likely to be prevalent among mushroom-forming Agaricomycetes and possibly unique to these fungi.

**Table 1.**
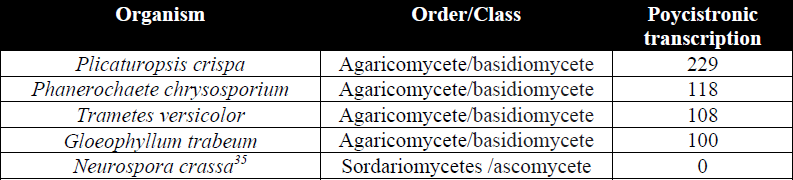
Polycistronic transcripts identified in several fungi transcriptomes

## DISCUSSION

Here we present a combined experimental and bioinformatics strategy (ToFU) that uses PacBio long reads for transcript isoform discovery. This strategy does not rely on reference genomes, and thereby enables the study of the transcriptome of any species. We showed that our strategy accurately reconstructs complex transcriptomes without relying on short-reads for error-correction or short-read assembly.

Lower mRNA isoform diversity has been observed in fungi compared to plants or animals. The low estimation is likely reflective of biased sampling from fungal lineages, such as the ascomycetes that have less complex gene structures and may have lower levels of isoform diversity^36^. The proficiency of ToFU was demonstrated on the transcriptome of the wood-degrading basidiomycete *P. crispa.* Our study shows that more than half of the genes in *P. crispa*, produce more than one transcript isoform, suggesting transcript isoform diversity in this phyla has likely been underestimated previously due to lack of deep full-length cDNA data. Similar to other non-fungal systems, genes producing the largest numbers of distinct isoforms are probable targets for regulation by NMD^22^. In this way alternative splicing may control production of functional proteins. Sequence optimization of GH and related enzymes may therefore be important in order to influence splicing and maximize production of transcript isoforms encoding functional enzymes in bioengineered systems.

A surprising finding is the discovery of long polycistronic transcripts spanning multiple independently transcribed loci that retain coding potential. Analysis of gene configuration conservation and long-read sequencing of multiple transcriptomes suggests that polycistronic transcripts may be found in mushroom-forming Agaricomycete fungi and absent from other basidiomycetes and ascomycetes. Future long-read transcriptome studies will resolve further details about the evolutionary breadth and origin of polycistronic readthrough transcription, as well as selective forces acting on the associated gene configurations. Interestingly, multi-ORF readthrough transcripts were associated with half of the bioinformatically detected secondary metabolite gene clusters identified by antiSMASH^37^. In the context of secondary metabolite gene clusters co-regulation and co-segregation can prevent the accumulation of toxic intermediates from these pathways^38^. Thus, polycistronic transcription may play an important role in achieving a particular ratio of enzymes produced from biosynthetic gene clusters, representing an advantageous mechanism to coordinate cellular responses.

A full understanding of the roles of polycistronic transcripts requires further experimental characterization of these polycistronic mRNAs. For example, how are they translated? Are they post-transcriptionally cleaved and processed? Post-transcriptional processing in response to environmental conditions has been shown for specific cases in other systems to regulate protein expression^40^. Conducting heterologous expression and biochemical characterization of the products encoded by some of these polycistronic mRNAs is a crucial step to understand the function and role of polycistronic transcripts in fungi. Manipulation and engineering of polycistronic transcripts has the potential to positively impact the field of bioconversion.

## METHODS

### Library preparation and cDNA sequencing

Total RNA was isolated from a monokaryotic culture of *Plicaturopsis crispa* grown on either a glucose-rich medium or microcrystalline cellulose. Total RNA was isolated from a monokaryotic cultures of *Phanerochaete chrysosporium,Trametes versicolor*, and *Gloeophyllum trabeum.* PolyA^+^ RNA was purified from total RNA via oligo-dT magnetic beads (Dynal). Four sequencing libraries (1-2 kb, 2-3 kb, 6kb, and no size-selection) were made for both P. crispa growth conditions and were prepared according to the PacBio isoform-sequencing protocol (http://www.smrtcommunity.com/servlet/servlet.FileDownload?file=00P7000000Pb1fkEAB). *Phanerochaete chrysosporium, Trametes versicolor*, and *Gloeophyllum trabeum* long-reads were generated from a size selected (>2 kb) cDNA library for each species. Single-molecule sequencing was performed on the PacBio RS II using P4-C2 chemistry, MagBead loading and 2 hour movie times.

### The ToFU pipeline

The pipeline consists of three stages: identifying full-length reads, isoform-level clustering, and final consensus polishing.

In the first stage, ToFU classifies all input raw reads into Circular Consensus Sequences (CCS) and non-CCS subreads by searching for the presence of sequencing adapters. Then ToFU determines a CCS or subread sequence to be full-length if both the 5’ and 3’ cDNA primers were present and there was a polyA tail signal preceding the 3’ primer.

In the second stage, ToFU uses an iterative isoform-clustering algorithm to cluster all the full-length reads derived from the same isoform. Briefly it first does clique-finding based on a similarity graph, then calls consensus using the Directed Acyclic Graph Consensus method and finally reassign sequences to different clusters based on their likelihood.

In the final stage, ToFU recruits the non-full-length reads and uses them to polish the consensus sequences produced during the second stage using the Quiver algorithm. Consensus sequences predicted to contain more than 10 errors are discarded.

### Merging the redundant PacBio transcripts into the ToFU transcript set

Due to the limitation of the cDNA library protocol, some cDNAs may not be full-length as they may lack the 5’-end. We collapsed transcripts that only differ in the 5’ start of their first exon but are otherwise identical in all subsequent exon structures keeping only the longest ones. The consequence of this step is that some transcripts with alternative transcription start sites are lost, but those with alternative splicing and alternatively polyadenylation will be preserved. This step can be avoided if the cDNA library protocol guarantees transcript sequences that preserve the 5’ start.

### Identification of polycistronic readthrough transcripts

We used Transdecoder for ORF prediction^41^. Transcripts with two or more non-overlapping ORFs ≥ 100 aa were further categorized based on reference annotations.

### RT-PCR and sequencing validation of the polycistronic RNAs

We selected 10 randomly selected polycistronic RNAs for experimental validation. RT-PCR primers were designed so that the target region begins near the end of the first ORF and ends within the second ORF. RT-PCR products were pooled and sequenced by PacBio sequencing. 29,511 raw reads were aligned to the 10 reference transcripts using BLASR. Only high quality end-to-end alignments (19,706 reads) were further analyzed. Eight out of 10 RT-PCR products exactly matched the references and therefore validated the polycistronic RNAs. The remaining two were inclusive, as one (scaffold_9:1201061-1204786) did not yield any matching sequencing reads, while the other (scaffold_15:638864-642834) had a different 3’ end from the designated 3’ target site. These two may represent RT-PCR off-target cases. Further details are listed in Supplemental Table 4.

### Poly-adenylation site (PAS) analysis

The poly-adenylation sites (PAS) of non-polycistronic and the second ORF of the polycistronic transcripts were identified by the polyA tail. The PAS of the first ORF of the polycistronic transcripts were identified with the aid of independent transcripts of the first ORF. The PAS motifs were predicted as previously described^30^.

### Short-read transcript reconstruction

PolyA^+^ RNA was purified from the same total RNA samples as used for long-read sequencing. 100-bp paired end Illumina reads were generated on the HiSeq2000 according to the manufacturer’s instructions (Illumina). Short-reads were assembled using Rnnotator (v.3.0.0), Oases (v0.2.08), and Cufflinks (v.2.1.1). Rnnotator and Oases are both *de novo* transcript assemblers whereas Cufflinks is reference-based. In order to obtain optimal assembly results for Oases, we performed eight assemblies with Oases using different values of k-mer then used Vmatch (v2.2.0) to remove redundancy. The k-mer size ranged from 53 to 95 and the step size was 6. For Cufflinks, short-reads were first aligned to the reference genome with TopHat (v2.0.6) then the alignments were assembled into a parsimonious set of transcripts using Cufflinks. All three programs were run with default options with strand-specific information.

Assembled transcripts were mapped to the reference genome using GMAP (v2014-04-24) using parameters ‘–cross-species–allow-close-indels 0–n 0’ and filtered for ≥ 99% alignment coverage and ≥ 85% alignment identity; these parameters are the same as those applied to the PacBio consensus sequences. Finally, the same redundancy removal script used for collapsing PacBio consensus sequences was applied to create a non-redundant, high-quality transcript set for each assembly program.

### Conservation of homologous gene configurations of polycistronic-associated gene pairs in other sequenced fungi

For identification of cases of gene order conservation of polycistronic gene pairs in other fungi (Supplemental Table S5), we searched for directly adjacent same-strand Blastp best hits in every fungal genome, publicly available at Mycocosm portal.

## Author Contributions

ZW and FC designed the study. ET developed the ToFU software. SG, ET, AS, JZ, DK and XM performed data analysis. ZZ and JU performed the experiments. SG, ET, IG, MF, JS and ZW wrote the paper.

## Acknowledgments

The authors thank Drs David Hibbett, Michele Weber and Ms. Sarah Middleton for their stimulating discussions and critical comments. The P. Crispa RNA samples were generously provided by Dr. David Hibbett. The work was conducted by the U.S. Department of Energy Joint Genome Institute and supported by the Office of Science of the U.S. Department of Energy under Contract No. DE-AC02-05CH11231.

## Data and Software Availability

The ToFU pipeline is released under the Standard PacBio Open Source License and has become an integrated module for the PacBio SMRTAnalysis tool suite (version 2.2 and up). The standalone version is available at: https://github.com/PacificBiosciences/cDNA_primer. All RNA-Seq data have been submitted to NCBI, under the BioProject ID: PRJNA261247.

## Supplementary Text

### 1. TOFU: a bioinformatics pipeline for PacBio transcriptome data

We developed a novel bioinformatics pipeline called TOFU to leverage both CCS (Circular Consensus Sequence) reads and non-CCS reads for transcript discovery. TOFU consists of three components: identifying full-length reads, isoform-level clustering, and final consensus polishing. We explain details in each step in the subsections below.

#### 1.1 Full-length read identification and artifact removal

Given either CCS or subreads, we use the *phmmer* program from HMMER^1^ to detect and remove the Clontech 5’/3’ primers (5’ – AAGCAGTGGTATCAACGCAGAGTAC – 3’). A read is considered full-length if both primers are detected at the ends with a polyA tail signal of at least 12 consecutive ‘A’s preceding the 3’ primer. Based on polyA tail and 3’ primer orientation, primer-trimmed reads are reverse complemented to represent the sense strand. Because the Clontech protocol does not ensure the capture of the 5’ cap, reads are considered 3’-complete but potentially 5’-partial; the 5’ incompleteness is taken into account in later stages of transcript collapsing. To remove artificial concatemers that may have formed via ligation of primer-attached inserts, the same *phmmer* program is used to detect the presence of Clontech primers at least 100 bp away from either end of the sequence.

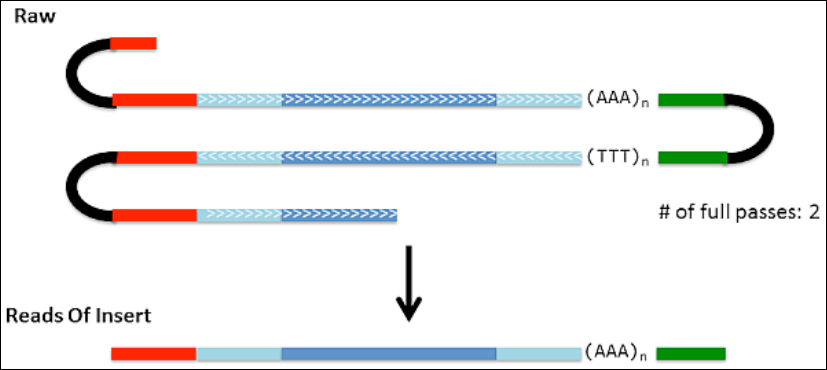

#### 1.2 Iterative isoform clustering & consensus calling using Quiver

We develop an iterative isoform clustering algorithm called ICE (Iterative Clustering for Error Correction) that uses PacBio sequencing QVs for determining whether two reads come from the same isoform. ICE consists of several main modules: (1) clique-finding based on similarity graph; (2) fast consensus calling with no QV information using DAGCon; (3) reassignment of sequences to different clusters based on likelihood. The following flow chart shows the process.

In the initial phase of clustering, the input sequence, which are often only a portion of the entire dataset, are aligned against each other using BLASR^2^ to construct a similarity graph where each node represents a read and each connecting edge indicates an “isoform hit”. Since BLASR is designed to align through long stretches of gaps, a hit between two transcripts that share some number of exons may have an alignment. To distinguish alignments from the same isoform while accounting for sequencing errors, an alignment between two reads is considered an “isoform hit” (from the same isoform) only if the percentage of gaps that cannot be attributed to base errors within a window size *w* is below some threshold *T.*

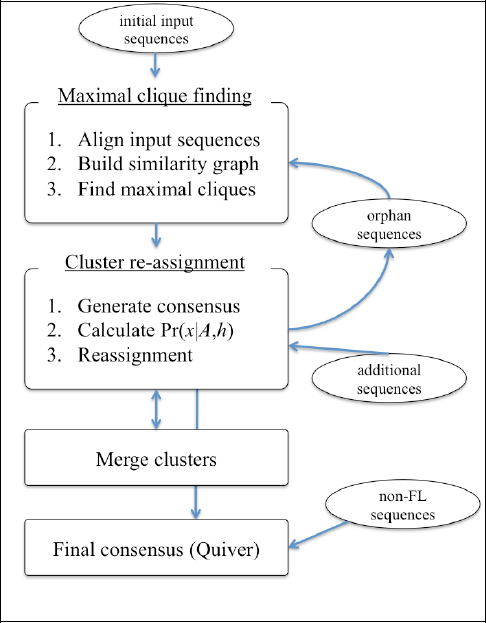

Formally, let the alignment string between two fully aligned sequences *x* and *y* be *A* = *a*_*1*_*a*_*2*_…*a*_*n*_ where *a*_*i*_, is ‘M’ for a match, ‘S’ for a substitution, ‘I’ for insertion, and ‘D’ for deletion (hence A is just an unraveled cigar string). Let 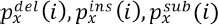 denote the probability of each error type based on the raw QVs for sequence *x*. Construct an non-match vector *E* = *e*_*1*_*e*_*2*_…*e*_*n*_ where *e*_*i*_ = 0 if one of the following is true:

- *a*_*i*_ = ‘M’
- *a*_*i*_ = ‘S’ and 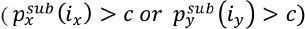
- *a*_*i*_ = ‘I’ and 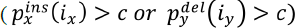
- *a*_*i*_ = ‘D’ and 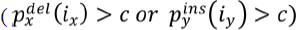

otherwise *e*_*i*_ = 1 which indicates a likely genuine non-match. Finally, we define *x* and *y* as being different isoforms if exists *i*, *j* > 0, where *j* − *i* < *w*, such that 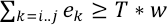. In other words, we identify indel-rich regions in the alignment that are likely due to exon-level differences. We use a previously published linear time algorithm for identifying indel-rich regions^3^. A pair of aligned sequences *x*, *y*, that do not have an indel-rich region, is considered an “isoform hit”. In this study, we set *c* = 0.1, *w* = 20, and *T* = 0.5.

With the similarity graph constructed using isoform hits, we look for perfect cliques in the graph. Ideally, all sequences from the same isoform would form a clique on its own with no other connecting edges. In practice, however, it is more likely that the sequences would form several cliques and may contain false positives (sequences from other isoforms). We address this by allowing “reassignment” of sequences to other clusters in a later step. For now, we run a maximal clique finding algorithm^4,5^ that non-deterministically finds maximal cliques in a graph, removes the clique nodes from the graph, then repeat the process until the entire graph is partitioned into mutually exclusive cliques (clusters).

We call an initial consensus on all clusters using DAGCon, a directed acyclic graph based consensus calling algorithm originally developed for error correcting PacBio genomic sequences^6^. With the improved accuracy of the consensus sequences, we can better approximate the likelihood of sequences belonging to the same isoform. Here, we use a “reassignment” procedure similar to the Gibbs sampling method described for detecting HIV quasispecies^7^. Briefly, we calculate the posterior probability of a sequence *x* originating from an isoform *h*(*c*) for cluster *c* as:

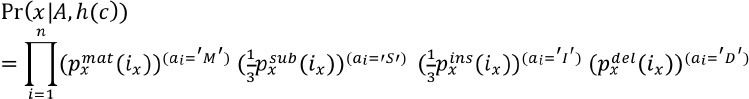

Theoretically, we need to calculate Pr(*x*|*A*, *h*(*c*)) for all sequences *x* and all cluster consensus *h*(*c*) ∈ **h**. In practice, only pairs of (*x*, *h*(*c*)), for which there is an “isoform hit” are calculated. Here “isoform hit” uses the same linear time algorithm in the similarity graph construction; the only difference is *h*(*c*) is considered error-less.

At each “reassignment” step, for each sequence *x*, there are three possible moves:

- Case 1: If no isoform hit exists for *x*, it is put into an “orphan” bin
- Case 2a: If there exists another cluster *c*’ such that Pr(*x*|*A*, *h*(*c*’)) > Pr(*x*|*A*, *h*(*c*)), reassign *x* to *c*’.
- Case 2b: If *x* is in a singleton cluster and there exists another cluster *c*’ such that *h*(*c*’) and *x* has an isoform hit, with some probability *p*, reassign *x* to *c*’.
- Case 3: If none of the above is true, *x* remains in *c*.

Case 2a deals with clusters that are big enough (>=3 sequences) to generate consensus. In cases where DAGCon cannot generate consensus because there is only 1 or 2 sequences in the cluster (called “singleton clusters”), Pr(*x*|*A*, *h*(*c*)) will always have the best probability and *x* will not have any possible moves. To allow the singletons to “escape”, we reassign it to another cluster for which there is an isoform hit with some low probability (by default, *p*=0.3).

At the end of the reassignments, the “orphan” sequences go through the same similarity graph construction and maximal clique finding process to form new clusters. Any cluster that has membership changes must have run through DAGCon again for consensus calling, as well as Pr(*x*|*A*, *h*) recalculated.

Because our algorithm does not jointly optimize for a global objective function (such as Pr (***h*| *A*, *x***), the total probability of observing the clusters given the input sequences and alignments) and our maximal clique finding is not guaranteed to put all isoform sequences in one cluster, a single isoform can end up being represented by multiple clusters. Thus, we add a phase of cluster merging, where the consensus sequences of two clusters are aligned against each other and if they are highly identical (≥ 99% similar) and are considered an isoform hit, then the two clusters are merged together. Note that, if two clusters were incorrectly merged, most commonly DAGCon will call consensus on one isoform but not the other, and as a result sequences belonging to the other isoform will be “orphaned” out in the next reassignment phase.

The iterative nature of the clustering process described so far makes adding new sequences very easy. New sequences can be introduced as follows: First, all new sequences are aligned against existing cluster consensus and assigned to the cluster with highest probability. For all sequences that did not have an isoform hit to an existing cluster, it is “orphaned” and follows the maximal clique finding procedure to be introduced into the dataset.

To summarize, the iterative process consists of maximal clique finding, consensus calling using DAGCon, cluster reassignment, and cluster merging. After a burn-in phase of reassignment and merging, the final set of DAGCon-generated consensus sequences are sent to the final stage of consensus calling using the more accurate and slower Quiver.

In this final stage of Quiver consensus calling, non-full-length reads, that were excluded from the iterative clustering process, is recruited to improve consensus accuracy. Non-full-length reads are aligned to all DAGCon-generated consensus sequences and filtered so that only “isoform hits” (using the same criterion as before but allowing for partial alignment) remain in the final alignment. Quiver uses the raw QVs from all aligned PacBio reads and outputs informative QVs along with the consensus sequence. Using the consensus QVs, we can filter out low quality consensus sequences that are often junk sequences and artifacts, though we also risk throwing out rare transcripts that have too little coverage.

Several speedups and parallelization are employed in the actual implementation of ICE. First, full-length reads are binned by size range (ex: 1-2k, 2-3k, 3-6k) since sequences from the same isoform must be within certain length differences even with indel errors. Partitioning the input sequences also serves to reduce the memory usage of each ICE process, which for efficiency maintains all QV information of “active” sequences (described below) in memory. Depending on the readlengths, 100k reads can take up 40-60GB of memory. Another speedup employed is to rerun DAGCon consensus calling only on clusters that are relatively small, where the removal of one or two sequences can affect the consensus sequence. In this study, we set the re-run cluster *A, h* does not to be re-calculated since it will remain the same. Finally, ICE maintains a set of “active” sequences that are sequences that are highly likely to be reassigned or orphaned because it is in a small cluster. A “freeze phase” is introduced after certain iterations of ICE, where any sequence that is in a cluster of size greater than the re-run size threshold and does not have an isoform hit to any other clusters is “inactivated” and forced to remain in its current cluster. QVs of inactive sequences are removed from memory. Consequently, clusters that contain inactive sequences, which must be of size greater than the re-run size threshold, does not ever have consensus re-run, and its core members can only increase.

#### 1.3 Software availability

As of this writing, TOFU has been incorporated into the official SMRTAnalysis suite (versions 2.2 and above) by Pacific Biosciences under the protocol name RS_IsoSeq. The only difference between TOFU and RS_IsoSeq is that while TOFU uses a mixture of CCS reads and subreads, RS_IsoSeq uses the improved ReadsOfInsert protocol that generates a consensus read for each ZMW. The developemental version of TOFU is available publicly at http://github.com/PacificBiosciences/cDNA_primer.

### 2. Short read mapping to long read consensus and filtering by coverage

Strand-specific Illumina paired short reads are treated as paired and concordantly mapped to the PacBio consensus sequences using BowTie2 with ‘–very-fast –norc’ and otherwise default parameters. Based on the short read coverage, PacBio consensus sequences are discarded if it: (1) has zero short read coverage that is not at the end of the sequence; or (2) has a sudden drop in coverage that is greater than 100X fold and the smaller coverage is less than 10.

### 3. Identifying exon and splice junctions and removing redundancy

PacBio consensus sequences are mapped to the *P. crispa* contigs using GMAP (version 2014-04-24) with parameters ‘–allow-close-indels 0 –cross-species’. Alignments with less than 99% coverage are discarded. Exon boundaries and alternative junctions are defined based on the remaining alignments. Because the PacBio reads are considered 3’ complete but possibly 5’ partial, transcripts are merged if they share the same 3’ exon and do not have any conflicting splice junctions. In the case of a single-exon transcript, all overlapping transcripts are merged.

### 4. ORF prediction and comparison with genome annotation

ORF prediction is done using TransDecoder^8^ on the PacBio (TOFU) transcript consensus sequences. To find polycistronic candidates, we filter for any PacBio transcripts that satisfy the following: (1) has two or more non-overlapping ORF predictions; (2) does not have another PacBio consensus with a single ORF prediction that maps to the same loci and whose predicted ORF length is between 80%–120% of the total ORF length (in aa) from (1). We ignore any polycistronic candidates that have a similar PacBio consensus with single predicted ORF because most of them appear to be either incompletely or faulty spliced. The filtered polycistronic transcripts are then categorized as either reference-supported or non-reference-supported depending on whether each of the predicted ORFs overlaps an annotated reference gene.

### 5. Identification of full-length CCS and subread sequences

We ran a total of 77 SMRTCells on the PacBio RS II consisting of different size fractions: 20 with no size selection, 22 from the 1-2k size selection, 19 from the 2-3k size selection, and 16 from the > 3k size selection. The loading efficiency (P1) for the runs were from 30-55%, which is the recommended range. The RS_Filter protocol from SMRTPortal (version 2.0) was used to generate filtered CCS (Circular Consensus Sequence) and subread sequences. Out of a total of 4,920,305 sequencing ZMWs, 1,628,297 (33%) were CCS ZMWs. We defined a CCS or subread sequence to be full-length (FL) if both the 5’ and 3’ cDNA primers were present and there was a polyA tail signal preceding the 3’ primer. Primers and polyA tails were trimmed from full-length sequences. Out of a total of 2,177,319 full-length CCS or non-CCS ZMWs, 4,748 were detected as artificial concatemers (0.2%) and were removed. The remaining trimmed, full-length sequences were further filtered for potential PCR chimeras by removing any sequence with at least 12 consecutive Ts in the beginning of the sequence. The remaining 2,548,103 sequences (from 2,143,039 ZMWs) were then used as input to the subsequent ICE clustering step.

### 6. Creating high-quality transcript consensus sequences

To speed up the clustering step, input sequences were divided into several bins (one for sequences shorter than 1kb, four for sequences between 1-2kb, two for sequences between 2-3kb, and one for sequences longer than 3kb) and ran through ICE independently on each bin. This resulted in many redundant transcripts consensus sequences that were merged in later steps. After obtaining the Quiver consensus sequence for each cluster, we estimated the number of expected errors based on the consensus QVs and discarded any consensus sequence that had more than 10 expected errors. While we risk throwing away rarer transcripts that have less coverage and thus worse consensus QVs, this ensured that the resulting consensus sequences were high quality. 40% of the clusters (176,903/443,242) passed this filter, which together consisted of 84% of the full-length input sequences. Most of the discarded clusters consisted of only one subread sequence, suggesting that these were likely low-quality sequences.

The high-quality consensus sequences were mapped to the *P. crispa* genome scaffolds using GMAP (version 2013-07-20) and removed for any sequence that did not map to the genome with at least 99% coverage; 12,085/179,603, or 6.8%, were removed.

### 7. Further filtering of PacBio consensus sequences based on short read evidence

Paired-end Illumina reads were mapped to the PacBio consensus sequences using BowTie2. Most of the short reads mapped to at least one PacBio consensus sequence. We found that most PCR chimeras that have formed during the full-length cDNA library construction in the PacBio reads were successfully filtered by the detection for polyT stretches in the sequence filtering steps. To exclude remaining PCR chimeras, we discarded any consensus sequence that did not have sufficient Illumina short read coverage throughout the sequence. We removed 17,335 out of 164,818 consensus sequences at this step. The remaining 147,483 consensus sequences then constitutes the redundant, high-quality transcript sequences that are supported by three independent sources: PacBio raw read support, Illumina short read support, and good alignment to the reference genome.

### 8. Collapsing redundant PacBio transcripts

Because the PacBio reads were considered 3’ complete but possibly 5’ partial, transcripts were merged if they shared the same 3’ exon and did not have any conflicting splice junctions. After merging redundant transcripts, we obtained 22,956 non-redundant transcript sequences in 9,073 isoform clusters.

### 9. Categorizing polycistronic readthrough transcripts

We screened for PacBio transcripts that had two or more non-overlapping ORF predictions that were not covered by another transcript that had a single, long ORF prediction. We then collapsed the readthrough transcripts by their mapped genomic locations and found 508 distinct loci to be polycistronic, among which 314 have support from genome-based gene predictions. The polycistronic transcripts were distributed across the genomic scaffolds and ranged from 828 to 5080 bp with an average length of 2330 bp. The majority of the candidates were bi-cistronic (471/508, or 93%), with the mean ORF lengths for the first and second ORFs being 256 aa and 277 aa .The mean distance between the two ORFs was 364 bp.

### Supplementary Tables

**Table S1.**
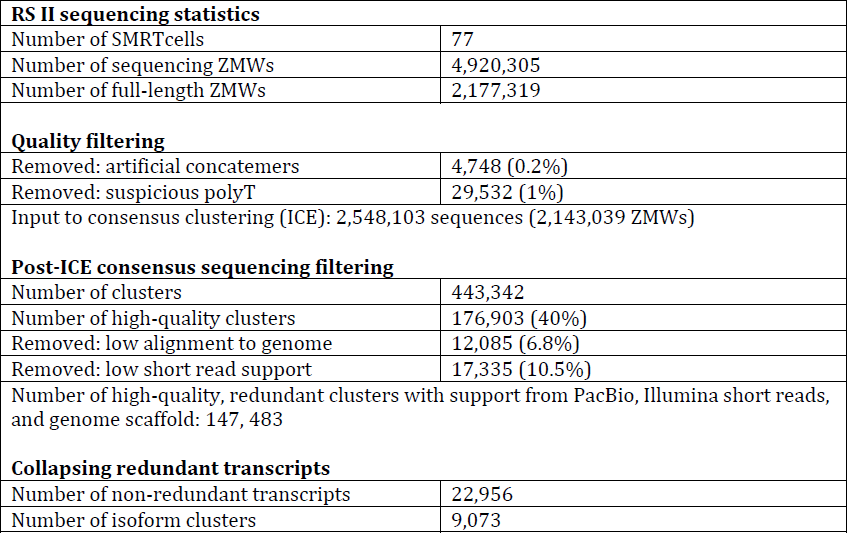
PacBio Sequencing statistics.

**Table S2.** Statistics for assembled transcripts from short reads.

| Program | Number of assembled transcripts | Well-mapped transcripts | Number of Non-redundant transcripts | Length of non-redundant transcripts (nt) |  |  |
| --- | --- | --- | --- | --- | --- | --- |
|  |  |  |  | Min | Max | Median |
| Rnnotator | 29,754 | 27,549 | 24,637 | 43 | 15,374 | 614 |
| Oases | 112,669 | 80,761 | 68,693 | 100 | 19,699 | 1,481 |
| Cufflinks | 10,211 | 10,184 | 10,051 | 96 | 20,688 | 1,960 |

**Table S3.** Comparison of assembled transcripts from short reads against PacBio transcripts.

| Program | Number of non-redundant transcripts | Match against PacBio* |  |  |  |  |  |
| --- | --- | --- | --- | --- | --- | --- | --- |
|  |  | Exact | Extended | Subset | Concordant | Alternative | Nomatch |
| Rnnotator | 24,637 | 3975 (16%) | 2488 (10%) | 5770 (23%) | 824 (3%) | 5171 (21%) | 6409 (26%) |
| Oases | 68,693 | 5212 (17%) | 20419 (33%) | 14351 (6%) | 4581 (3%) | 11081 (22%) | 13049 (19%) |
| Cufflinks | 10,051 | 1684 (8%) | 3308 (30%) | 590 (21%) | 294 (7%) | 2220 (16%) | 1955 (19%) |

\* Each assembled transcript was matched against PacBio transcripts and categorized based on the number and exact position of donor-acceptor sites, regardless of the start and end position of the first and last exon. An exact match indicates that the exon junctions are the same, whereas extended, subset, and concordant indicates exon junction agreement where there is overlap but there are additional junctions not covered by either the assembled transcript or PacBio. An alternative match means disagreement in exon junctions but the mapped loci overlaps. Finally, a nomatch indicates no PacBio is observed at the loci.

**Table S4.**
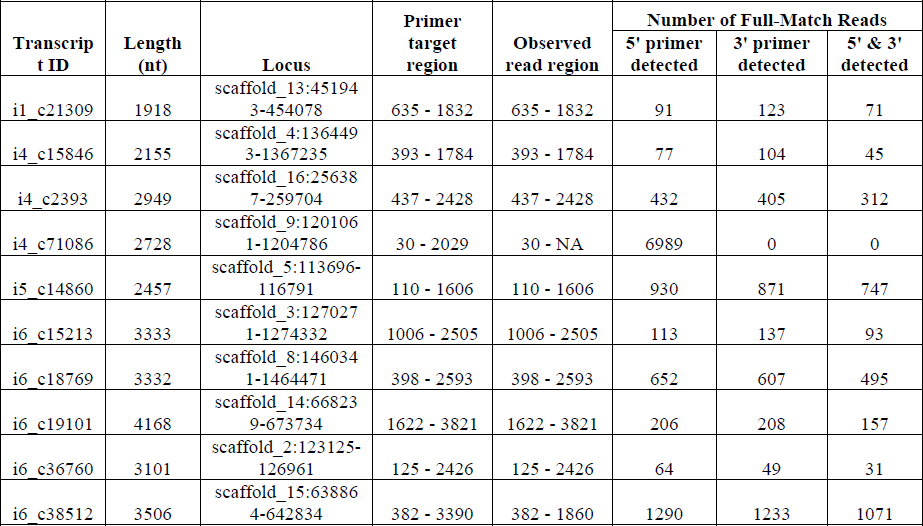
RT-PCR validation of polycistronic transcripts. Primers were designed to cover more than one of the predicted ORF regions to prevent mis-validation by sequencing of non-polycistronic transcripts that contain only one of the ORFs. Eight out of ten of the RT-PCR products and subsequent sequencing confirmed the presence of the polycistronic transcripts.

**Table S5.**
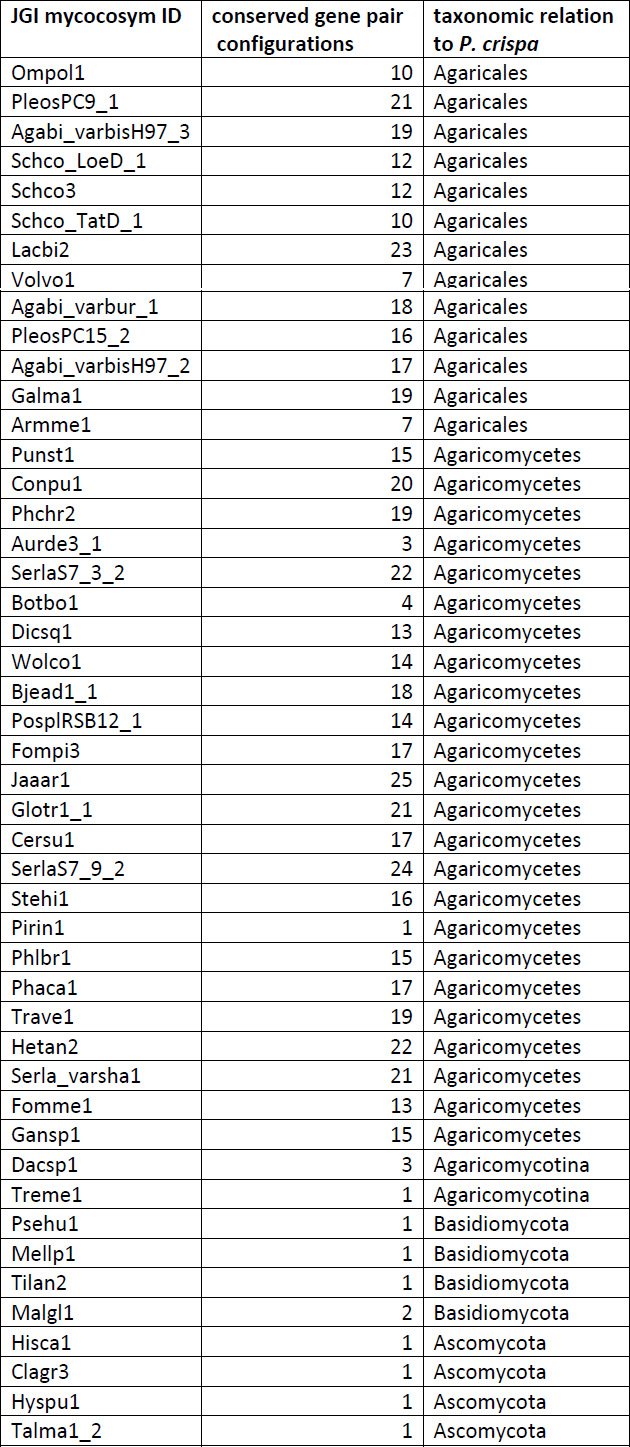
The list of species that are used for searching conserved gene pairs.

**Figure S1.**
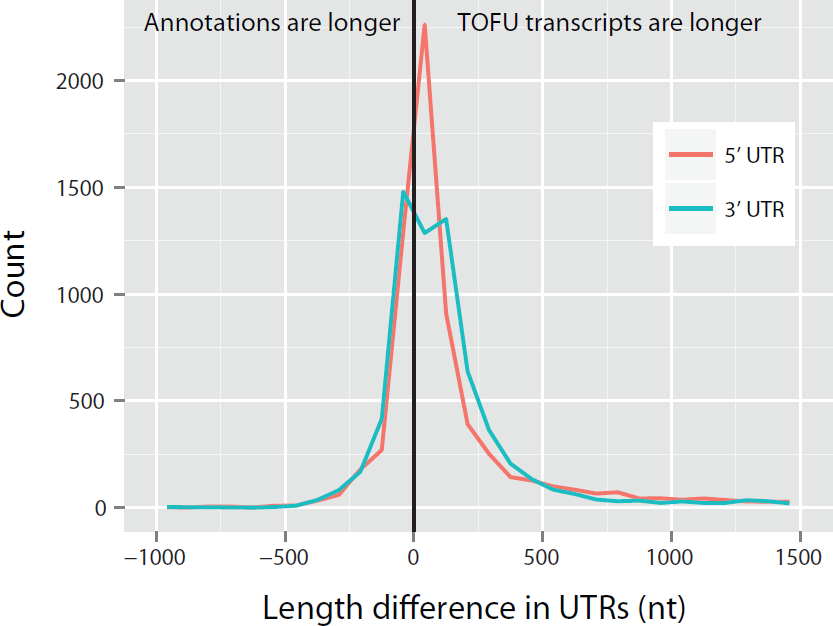
Most of the TOFU transcripts have longer UTRs than current annotation.

**Figure S2.**
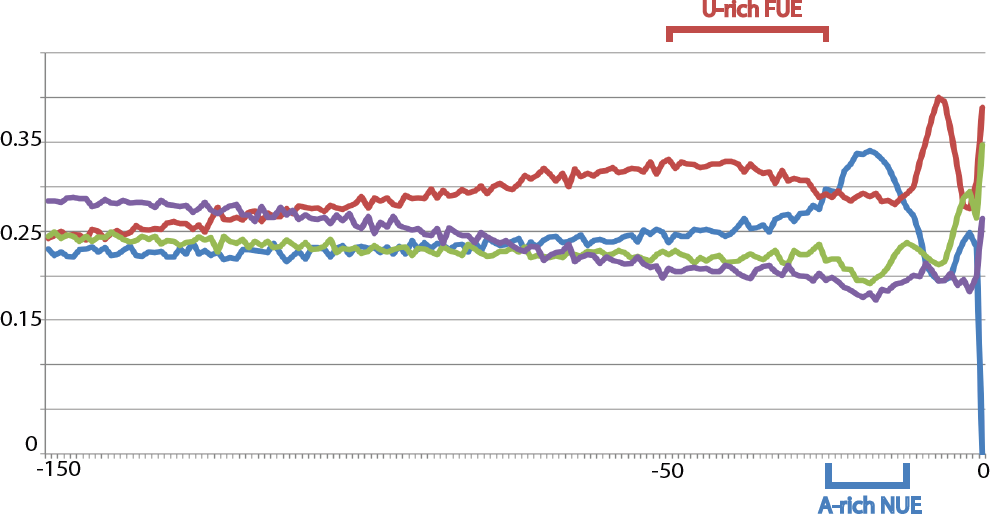
Genome-wide analysis of the transcription termination signals in P. crispa. Average nucleotide composition was plotted for all non-polycistronic ToFU transcripts upstream of the poly-adenylation site. A-rich NUE and U-rich FUE elements are indicated. Green represents C-content and purple represent G-content, respectively. X-axis indicates the distance to polyA site.

## References

1. Floudas, D. et al. The Paleozoic origin of enzymatic lignin decomposition reconstructed from 31 fungal genomes. Science 336, 1715–1719 (2012).

2. Ulitsky, I. & Bartel, D.P. lincRNAs: genomics, evolution, and mechanisms. Cell 154, 26–46 (2013).

3. Kung, J.T., Colognori, D. & Lee, J.T. Long noncoding RNAs: past, present, and future. Genetics 193, 651–669 (2013).

4. Matlin, A.J., Clark, F. & Smith, C.W. Understanding alternative splicing: towards a cellular code. Nat Rev Mol Cell Biol 6, 386–398 (2005).

5. Proudfoot, N.J. Ending the message: poly(A) signals then and now. Genes Dev 25, 1770–1782 (2011).

6. Di Giammartino, D.C., Nishida, K. & Manley, J.L. Mechanisms and consequences of alternative polyadenylation. Mol Cell 43, 853–866 (2011).

7. Parra, G. et al. Tandem chimerism as a means to increase protein complexity in the human genome. Genome Res 16, 37–44 (2006).

8. Akiva, P. et al. Transcription-mediated gene fusion in the human genome. Genome Res 16, 30–36 (2006).

9. Munk, C. et al. Functions, structure, and read-through alternative splicing of feline APOBEC3 genes. Genome Biol 9, R48 (2008).

10. Nagalakshmi, U. et al. The transcriptional landscape of the yeast genome defined by RNA sequencing. Science 320, 1344–1349 (2008).

11. Mortazavi, A., Williams, B.A., McCue, K., Schaeffer, L. & Wold, B. Mapping and quantifying mammalian transcriptomes by RNA-Seq. Nat Methods 5, 621–628 (2008).

12. Martin, J.A. & Wang, Z. Next-generation transcriptome assembly. Nat Rev Genet 12, 671–682 (2011).

13. Treutlein, B., Gokce, O., Quake, S.R. & Sudhof, T.C. Cartography of neurexin alternative splicing mapped by single-molecule long-read mRNA sequencing. Proc Natl Acad Sci U S A 111, E1291–1299 (2014).

14. Shearwin, K.E., Callen, B.P. & Egan, J.B. Transcriptional interference–crash course. Trends Genet 21, 339–345 (2005).

15. Thomas, S., Underwood, J.G., Tseng, E. & Holloway, A.K. Long-read sequencing of chicken transcripts and identification of new transcript isoforms. PLoS One 9, e94650 (2014).

16. Sharon, D., Tilgner, H., Grubert, F. & Snyder, M. A single-molecule long-read survey of the human transcriptome. Nat Biotechnol 31, 1009–1014 (2013).

17. Au, K.F. et al. Characterization of the human ESC transcriptome by hybrid sequencing. Proc Natl Acad Sci U S A 110, E4821–4830 (2013).

18. Koren, S. et al. Hybrid error correction and de novo assembly of singlemolecule sequencing reads. Nat Biotechnol 30, 693–700 (2012).

19. Grutzmann, K. et al. Fungal alternative splicing is associated with multicellular complexity and virulence: a genome-wide multi-species study. DNA Res 21, 27–39 (2014).

20. Grigoriev, I.V. et al. MycoCosm portal: gearing up for 1000 fungal genomes. Nucleic Acids Res 42, D699–704 (2014).

21. Wu, T.D. & Watanabe, C.K. GMAP: a genomic mapping and alignment program for mRNA and EST sequences. Bioinformatics 21, 1859–1875 (2005).

22. Filichkin, S.A. et al. Genome-wide mapping of alternative splicing in Arabidopsis thaliana. Genome Res 20, 45–58 (2010).

23. Wang, E.T. et al. Alternative isoform regulation in human tissue transcriptomes. Nature 456, 470–476 (2008).

24. Trapnell, C. et al. Differential analysis of gene regulation at transcript resolution with RNA-seq. Nat Biotechnol 31, 46–53 (2013).

25. Martin, J. et al. Rnnotator: an automated de novo transcriptome assembly pipeline from stranded RNA-Seq reads. BMC Genomics 11, 663 (2010).

26. Schulz, M.H., Zerbino, D.R., Vingron, M. & Birney, E. Oases: robust de novo RNA-seq assembly across the dynamic range of expression levels. Bioinformatics 28, 1086–1092 (2012).

27. Calvo, S.E., Pagliarini, D.J. & Mootha, V.K. Upstream open reading frames cause widespread reduction of protein expression and are polymorphic among humans. Proc Natl Acad Sci U S A 106, 7507–7512 (2009).

28. Blumenthal, T. Operons in eukaryotes. Brief Funct Genomic Proteomic 3, 199–211 (2004).

29. Shen, Y., Liu, Y., Liu, L., Liang, C. & Li, Q.Q. Unique features of nuclear mRNA poly(A) signals and alternative polyadenylation in Chlamydomonas reinhardtii. Genetics 179, 167–176 (2008).

30. Tanaka, M., Sakai, Y., Yamada, O., Shintani, T. & Gomi, K. In silico analysis of 3’-end-processing signals in Aspergillus oryzae using expressed sequence tags and genomic sequencing data. DNA Res 18, 189–200 (2011).

31. Lee, H.C. et al. Diverse pathways generate microRNA-like RNAs and Dicer-independent small interfering RNAs in fungi. Mol Cell 38, 803–814 (2010).

32. Gullerova, M. & Proudfoot, N.J. Cohesin complex promotes transcriptional termination between convergent genes in S. pombe. Cell 132, 983–995 (2008).

33. Hainer, S.J., Pruneski, J.A., Mitchell, R.D., Monteverde, R.M. & Martens, J.A. Intergenic transcription causes repression by directing nucleosome assembly. Genes Dev 25, 29–40 (2011).

34. Palmer, A.C., Ahlgren-Berg, A., Egan, J.B., Dodd, I.B. & Shearwin, K.E. Potent transcriptional interference by pausing of RNA polymerases over a downstream promoter. Mol Cell 34, 545–555 (2009).

35. Kim, K. et al. Long-read, whole-genome shotgun sequence data for five model organisms: E. coli, S. cerevisiae, N. crassa, A. thaliana, and D. melanogaster. (2014).

36. Loftus, B.J. et al. The genome of the basidiomycetous yeast and human pathogen Cryptococcus neoformans. Science 307, 1321–1324 (2005).

37. Blin, K. et al. antiSMASH 2.0–a versatile platform for genome mining of secondary metabolite producers. Nucleic Acids Res 41, W204–212 (2013).

38. McGary, K.L., Slot, J.C. & Rokas, A. Physical linkage of metabolic genes in fungi is an adaptation against the accumulation of toxic intermediate compounds. Proc Natl Acad Sci U S A 110, 11481–11486 (2013).

39. Dutartre, L., Hilliou, F. & Feyereisen, R. Phylogenomics of the benzoxazinoid biosynthetic pathway of Poaceae: gene duplications and origin of the Bx cluster. BMC Evol Biol 12, 64 (2012).

40. Prasanth, K.V. et al. Regulating gene expression through RNA nuclear retention. Cell 123, 249–263 (2005).

41. Haas, B.J. et al. De novo transcript sequence reconstruction from RNA-seq using the Trinity platform for reference generation and analysis. Nat Protoc 8, 1494–1512 (2013).

## References

1. Eddy, S.R. Accelerated Profile HMM Searches. PLoS Comput Biol 7, e1002195 (2011).

2. Chaisson, M.J. & Tesler, G. Mapping single molecule sequencing reads using basic local alignment with successive refinement (BLASR): application and theory. BMC Bioinformatics 13 (2012).

3. Tseng, H.H. & Tompa, M. Algorithms for locating extremely conserved elements in multiple sequence alignments. BMC Bioinformatics 10, 432 (2009).

4. Abello, J., P. M. Pardalos, and M. G. C Resende On Maximum Clique Problems in Very Large Graphs. AT&T Labs Reserrch Technical Report TR98 (1998).

5. Tseng, H.-H. Discovery and Applications of Bacterial Noncoding RNAs. PhD thesis, Department of Computer Science & Engineering: University of Washington (2012).

6. Chin, C.S. et al. Nonhybrid, finished microbial genome assemblies from long-read SMRT sequencing data. Nat Methods 10, 563–569 (2013).

7. Zagordi, O., Geyrhofer, L., Roth, V. & Beerenwinkel, N. Deep sequencing of a genetically heterogeneous sample: local haplotype reconstruction and read error correction. J Comput Biol 17, 417–428 (2010).

8. Haas, B.J. et al. De novo transcript sequence reconstruction from RNA-seq using the Trinity platform for reference generation and analysis. Nat Protoc 8, 1494–1512 (2013).

